# A member of the tryptophan-rich protein family is required for efficient sequestration of *Plasmodium berghei* schizonts

**DOI:** 10.1101/2022.07.14.500060

**Authors:** Julie-Anne Gabelich, Josephine Grützke, Florian Kirscht, Oliver Popp, Joachim M Matz, Gunnar Dittmar, Melanie Rug, Alyssa Ingmundson

## Abstract

Protein export and host membrane remodeling are crucial for multiple *Plasmodium* species to establish a niche in infected hosts. To better understand the contribution of these processes to successful parasite infection *in vivo*, we sought to find and characterize protein components of the intraerythrocytic *Plasmodium berghei*-induced membrane structures (IBIS) that form in the cytoplasm of infected erythrocytes. We identified proteins that immunoprecipitate with IBIS1, a signature member of the IBIS in *P. berghei*-infected erythrocytes. In parallel, we also report our data describing proteins that co-precipitate with the PTEX (*Plasmodium* translocon of exported proteins) component EXP2. To validate our findings, we examined the location of three candidate IBIS1-interactors that are conserved across multiple *Plasmodium* species, and we found they localized to IBIS in infected red blood cells and two further co-localized with IBIS1 in the liver-stage parasitophorous vacuole membrane. Successful gene deletion revealed that these two tryptophan-rich domain-containing proteins, termed here IPIS2 and IPIS3 (for intraerythrocytic *Plasmodium*-induced membrane structures), are required for efficient blood-stage growth. Erythrocytes infected with IPIS2-deficient schizonts in particular fail to bind CD36 as efficiently as wild-type *P. berghei*-infected cells and therefore fail to effectively sequester out of the circulating blood. Our findings support the idea that intra-erythrocytic membrane compartments are required across species for alterations of the host erythrocyte that facilitate interactions of infected cells with host tissues.

**Author Summary:** Red blood cells, which are typically devoid of organelles or other intracellular membrane compartments, are host to *Plasmodium* parasites in a malaria infection. These intracellular parasites export proteins into the host red blood cell cytoplasm and generate novel membranous organelles therein. The best characterized of these membrane structures are known as Maurer’s clefts in *Plasmodium falciparum*-infected cells; however, infection with any studied *Plasmodium* species leads to the generation of membrane structures in the host red blood cell. For these other *Plasmodium* species, the known protein repertoire of these cleft-like structures is extremely limited. Our study expands upon this repertoire in the rodent parasite *Plasmodium berghei*. We genetically targeted two of the proteins we identified in these cleft-like structures and found both are required for efficient *Plasmodium* growth in the host’s blood. One of these, which we term IPIS2, is required for the binding of late-stage *Plasmodium*-infected red blood cells to the vascular endothelium to sequester out of the circulating blood. Both proteins have a tryptophan-rich domain, and this is the first time a protein with this domain has been found to affect the remodeling of the host red blood cell during *Plasmodium* infection.

## Introduction

Well-tuned host-parasite interactions allow *Plasmodium* parasites to be effective pathogens, which results in more than half of the world’s population to be at risk of contracting malaria caused by one of the six *Plasmodium* species able to infect humans.

*Plasmodium* parasites successfully infect two cell types in their vertebrate hosts: the initial liver stage of infection is followed by the infection of blood cells, the stage causing disease. Malaria parasites strategically remodel the red blood cells in which they reside. These adaptations create an intracellular environment suitable for parasite development and generate vessels that deftly navigate the host organism. Host alterations are carried out by proteins delivered from the parasite into the host cell. These exported *Plasmodium* proteins are delivered across the parasitophorous vacuole membrane into the host cell via the parasite-encoded PTEX protein export apparatus [1-3]. The effects of protein export have been primarily described in *Plasmodium falciparum* where delivery and presentation of the *P. falciparum*-specific adhesion protein *Pf*EMP1 on the surface of infected red blood cells is a fundamental consequence of protein export. Protein export and host remodeling are not, however, exclusive to this *Plasmodium* species. The PTEX components are conserved across species [1], and infection of red blood cells with any described *Plasmodium* species leads to alterations of the host cell [4-8]. These alterations can include the transport of parasite proteins to the erythrocyte surface in addition to the morphologically evident membrane alterations [9-13].

The establishment of membrane compartments in the red blood cell cytoplasm of infected red blood cells are found across *Plasmodium* species. Although these structures have various morphologies and different names including Maurer’s clefts in *P. falciparum*-infected cells [7], Sinton Mulligan stipplings/clefts in *P. knowlesi*-infected cells [14], Schüffner’s dots or caveolae-vesicle complexes in *P. vivax*-infected cells [6], and intra-erythrocytic *P. berghei*-induced structures (IBIS) in *P. berghei*-infected cells [15, 16], the conservation of a few “cleft”-localized proteins suggests that at least some of the functions of these membrane compartments are conserved across species [17-19]. When genetic studies have targeted *P. falciparum* proteins that localize to Maurer’s clefts, the gene deletions affect presentation of PfEMP1 on the surface of the red blood cells, knob morphology and/or Maurer’s cleft morphology [20-27], which are all unique to *P. falciparum*-infected cells.

To learn more about the host cell-localized membrane modifications in other *Plasmodium* species we used the rodent malaria parasite, *P. berghei* as a model. We aimed to investigate the contribution of these conserved membrane features in a system in which we can investigate the consequences of targeting exported proteins on parasite success *in vivo*. To identify additional protein components that localize to these membrane compartments in *Plasmodium*-infected cells, we immunoprecipitated an IBIS signature protein, IBIS1, from *P. berghei*-infected red blood cells. We focused on two proteins that co-precipitated with IBIS1, PBANKA_0524300 and PBANKA_0623100, which we found localize to IBIS1-positive membranes in the liver and blood infection stages. We termed these proteins intra-erythrocytic *<u>P</u>lasmodium-*induced <u>s</u>tructure proteins 2 and 3 (IPIS2 and IPIS3, respectively) and found that while they are both required for robust parasite propagation in the blood, loss of IPIS2 specifically prevents efficient sequestration of *P. berghei* schizonts from the blood circulation.

## Results

### Identification of *Plasmodium* proteins co-precipitating with IBIS1-mCherry and EXP2-mCherry

To identify additional protein components of the IBIS membrane structures, we immunoprecipitated IBIS1-mCherry from infected erythrocytes. As controls, we also immunoprecipitated mCherry from erythrocytes infected with other *P. berghei* lines expressing mCherry or mCherry fusion proteins (Figure 1A). In these control samples, mCherry was localized in the cytoplasm of the parasite (labeled in figure 1a as mCherry) [28], the cytoplasm of the host erythrocyte (PEXEL-mCherry) [29], or in the parasitophorous vacuole membrane (EXP2-mCherry) [30]. In the EXP2-mCherry line, full length EXP2 is tagged with mCherry and expressed from the endogenous promoter. Because EXP2 is a well-characterized *Plasmodium* protein that is a component of the *Plasmodium* translocon of exported proteins (PTEX), immunoprecipitation of EXP2-mCherry further served as a positive control for the pulldown assay since it is known to interact with other PTEX components [31, 32].

**Figure 1.**
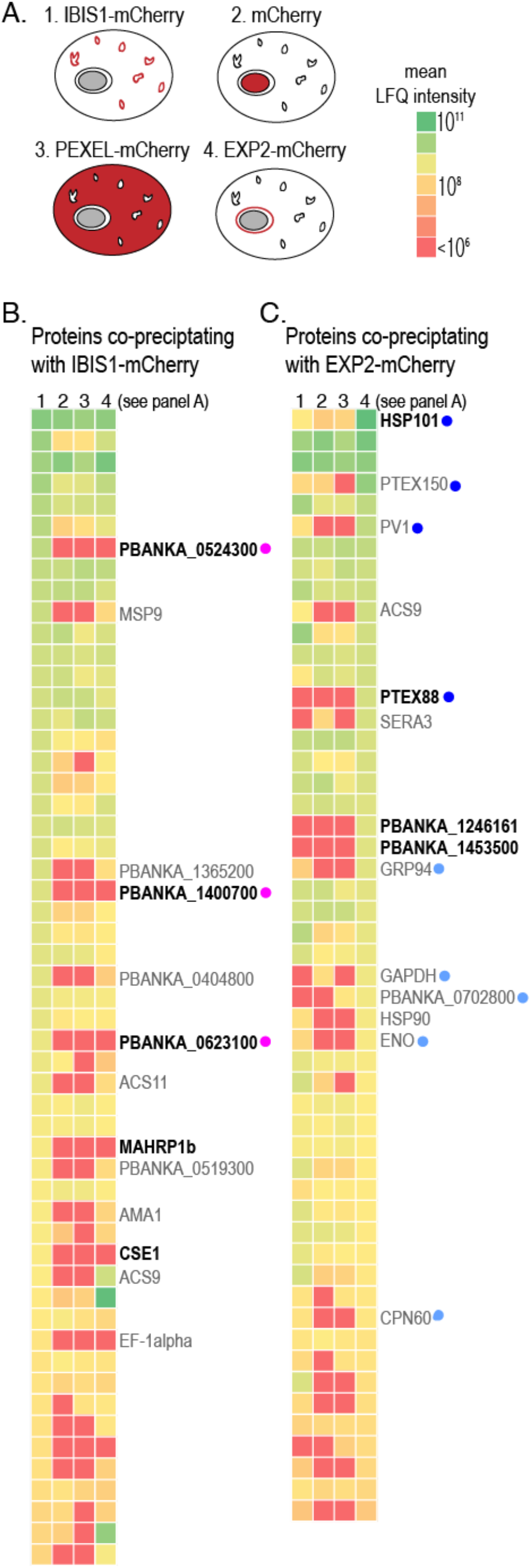
Immunoprecipitation of *P. berghei* proteins from infected red blood cells identifies proteins that co-precipitate specifically with IBIS1-mCherry and EXP2-mCherry. A) The indicated mCherry-tagged proteins were immunoprecipitated from lysates of late stage *P. berghei* using RFP-specific nanobodies and co-precipitating proteins were identified by LC-MS/MS. B) *P. berghei* proteins that were detected in the IBIS1-mCherry sample in at least two of the experimental replicates are depicted in the heatmap, which indicate the mean normalized LFQ intensities of detected proteins of three independent experiments. Each row represents a protein detected in the IBIS1-mCherry eluates. The columns represent the immunoprecipitates prepared from the mCherry-expressing parasite lines indicated in (A) (1: IBIS1-mCherry, 2: mCherry, 3: PEXEL-mCherry, 4:EXP2-mCherry). Proteins that were enriched in the IBIS1-mCherry pulldown relative to at least two of the three controls (top 10% of the normal distribution) are annotated. Those in bold were enriched relative to all three controls. Proteins marked with a pink dot were followed up further in this study. C) *P. berghei* proteins that were detected in the EXP2-mCherry sample in at least two of the experimental replicates are depicted in the heatmap, which indicate the mean normalized LFQ intensities of detected proteins of three independent experiments. Each row represents a protein detected in the EXP2-mCherry eluates. The columns represent the immunoprecipitates prepared from the different mCherry-expressing parasite lines indicated in (A). Proteins marked with dark blue dots are confirmed to play roles in PTEX-mediated export, and those marked with light blue dots were previously found biochemically associated with components of the PTEX complex [33, 34]

The mCherry-tagged proteins were immunoprecipitated with anti-RFP nanobody-coupled resin and precipitating proteins were analyzed by liquid chromatography – tandem mass spectrometry (LC-MS/MS). The results were analyzed based on the label-free quantitation (LFQ) intensities of the detected peptides (full results in Table S1). Differences in starting material quantities affected the overlap of peptides detected in all three independent experimental replicates. We therefore focused on *Plasmodium* proteins detected in the IBIS1-mCherry or EXP2-mCherry pulldown in at least two of the three independent experimental replicates, and analyzed enrichment using the mean LFQ intensities across all three replicates (Figure 1B).

Several of the established protein components of the PTEX complex [1], including HSP101, PTEX150 and PTEX88 co-precipitated specifically with EXP2-mCherry (Figure 1C). Several additional proteins that have previously co-purified with PTEX proteins [33, 34] were also identified in our analysis as being specifically enriched in the EXP2-mCherry sample. Furthermore, a protein previously identified in the intra-erythrocytic IBIS of *P. berghei*-infected cells, *Pb*MAHRP1 [17], co-precipitated specifically with IBIS1-mCherry. Together, these results confirm the specificity of the pulldowns and validity of the analyses.

### The tryptophan-rich proteins IPIS2 and IPIS3 colocalize with IBIS1 in both liver and blood stages of infection

Four proteins, in addition to *Pb*MAHRP1, were enriched in the IBIS1-mCherry immunoprecipitation relative to all three controls: PBANKA_0524300, PBANKA_0623100, PBANKA_1400700, and CSE1, which is annotated as a putative importin alpha. The three proteins with the highest abundance in the IBIS1-mCherry sample, PBANKA_0524300, PBANKA_0623100 and PBANKA_1400700, were selected for further analysis (Figure 2A). Two of the three proteins do not have predicted secretion or export signals, but do have a tryptophan-threonine rich domain (PF12319) that is conserved in several proteins across *Plasmodium* species. To visualize the expression and localization of these proteins, they were each C-terminally tagged with the fluorescent protein mCherry via integration of the tag at the endogenous gene locus (Figure S1A, B).

**Figure 2.**
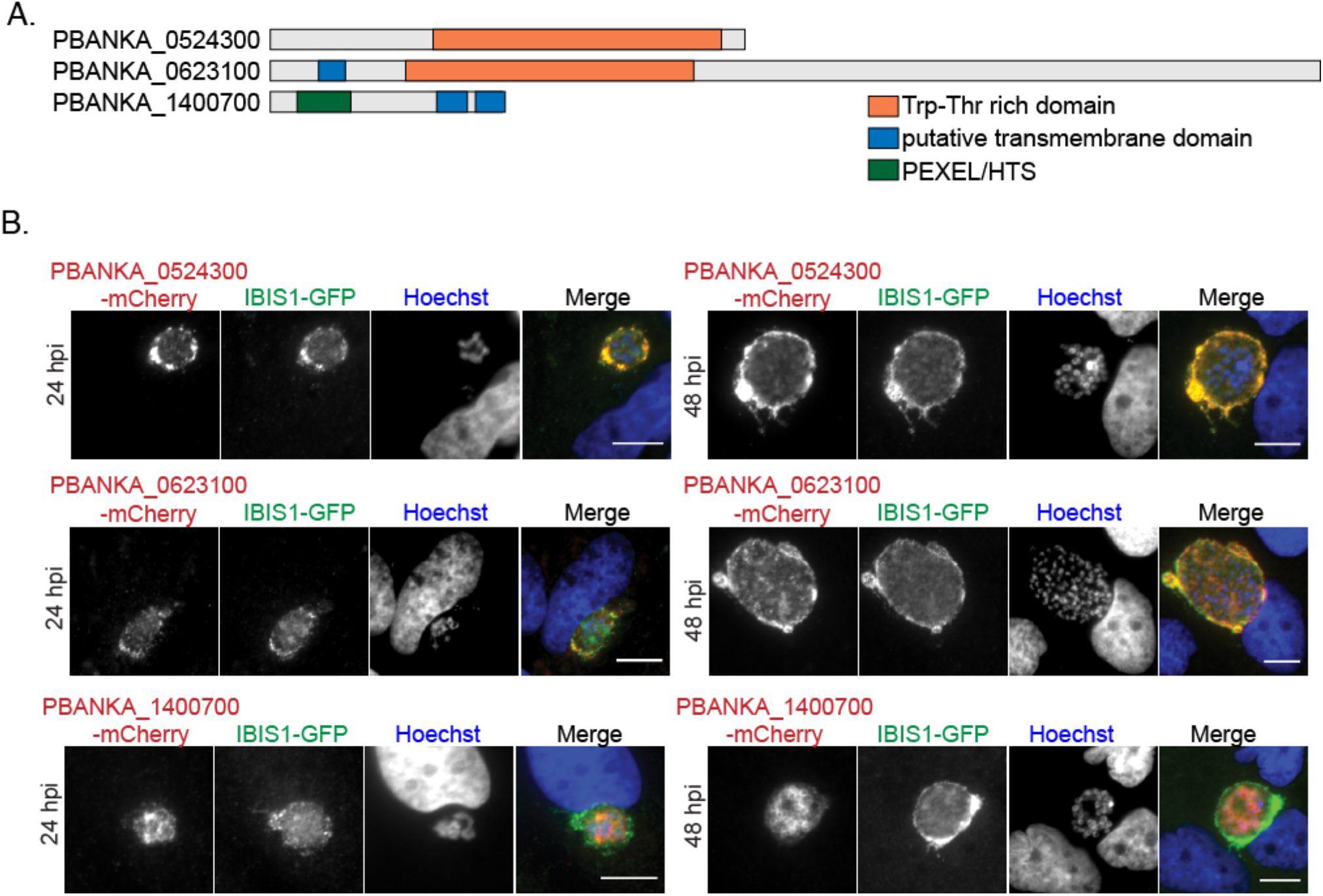
mCherry tagged PBANKA_0524300 (IPIS2), PBANKA_0623100 (IPIS3) colocalize with IBIS1-GFP at the liver-stage parasitophorous vacuole. A) Protein features of candidate proteins that coprecipitated with IBIS1. B) Hepatoma cells were infected with sporozoites after the cross-fertilization of IBIS1-GFP and either PBANKA_0524300-mCherry, PBANKA_0623100-mCherry or PBANKA_1400700-mCherry transgenic parasite lines. Cells were fixed at indicated time points and the mCherry and GFP signals were amplified by using anti-RFP and anti-GFP antibodies. Scale bars, 10 μm.

All three proteins were found to be exported to the erythrocyte cytoplasm (Figure S1C). Consistent with previous reports [11], PBANKA_0623100-mCherry localizes to punctate structures in the cytoplasm of infected red blood cells (Figure S1C). We found PBANKA_0524300-mCherry exhibits a similar punctate localization in the erythrocyte cytoplasm. A previous study reported exported GFP-tagged PBANKA_1400700 evenly distributed throughout the erythrocyte cytoplasm [35]. In contrast, we detected the exported fusion protein in a punctate pattern (Figure S1C). In both this study and the previous study, much of the fluorescently tagged PBANKA_1400700 remained associated with the parasites. While all three of the tagged candidate proteins could be seen localizing to the parasite in some infected cells, this localization was most pronounced for PBANKA_1400700-mCherry.

All three proteins were also detected in the liver infection stage (Figure S2A). PBANKA_0524300-mCherry and PBANKA_0623100-mCherry were first detected 24 hours post-infection in the developing liver-stage forms and surround the parasite in a pattern indicative of the liver-stage parasitophorous vacuole and associated tubovesicular network. PBANKA_1400700-mCherry was detected throughout the liver stage, but it was not secreted from the parasite despite the presence of a signal peptide.

To determine if the localization of these three proteins overlaps with IBIS1, the parasite lines in which these proteins are tagged with mCherry were crossed with a line in which IBIS1 is tagged with GFP via co-infections of mosquitoes. A proportion of the parasites resulting from the cross express both fluorescent proteins. Signals from PBANKA_0524300-mCherry and PBANKA_0623100-mCherry overlapped considerably with the IBIS1-GFP signal surrounding the liver-stage parasites (Figure 2B). This colocalization was confirmed with confocal microscopy (Figure S2B) and indicates that IBIS1, PBANKA_0524300 and PBANKA_0623100 are present within the same domains of the liver-stage parasitophorous vacuole and tubovesicular network. In infected red blood cells, PBANKA_0524300-mCherry and PBANKA_0623100-mCherry colocalize with IBIS1-GFP. Although the majority of PBANKA_1400700-mCherry remains parasite-associated, the exported mCherry signal colocalizes with IBIS1-GFP. These results indicate that all three studied proteins localize to the same membrane compartments formed in the infected erythrocyte cytoplasm (Figure 3A).

**Figure 3.**
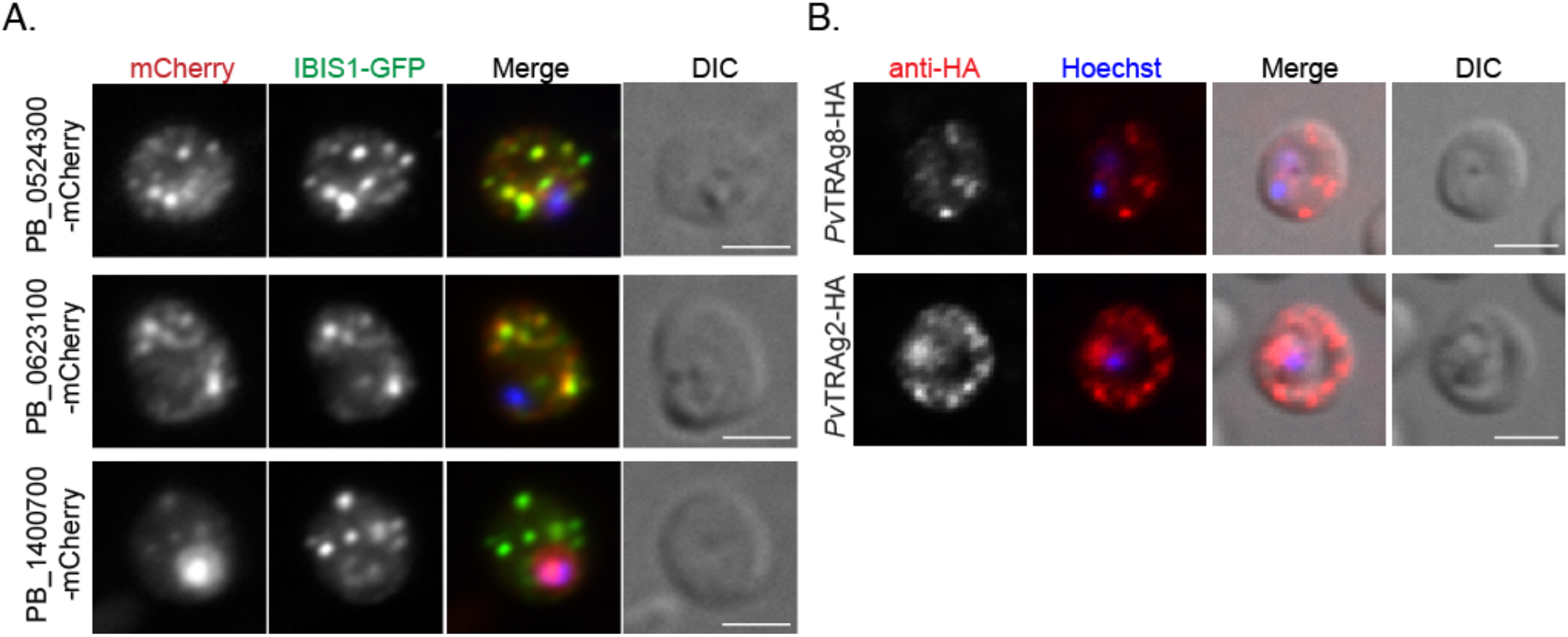
PBANKA_0524300 (IPIS2), PBANKA_0623100 (IPIS3), PBANKA_1400700 and putative *P. vivax* orthologs localize to discrete structures in the cytoplasm of infected erythrocytes. A) PBANKA_0524300-mCherry, PBANKA_0623100-mCherry, and PBANKA_1400700-mCherry co-localize with the IBIS marker IBIS1-GFP in the erythrocyte cytoplasm. Infected cells were fixed and labeled with anti-GFP and anti-mCherry antibodies to amplify the fluorescent signal. Scale bars, 5 μm. B) Erythrocytes infected with *P. berghei* expressing the HA-tagged *P. vivax* proteins *Pv*TRAg8 or *Pv*TRAg2, which are putative orthologs of IPIS2 and IPIS3, respectively, were fixed and labeled with anti-HA antibodies. Scale bars, 5 μm.

Because PBANKA_0524300 and PBANKA_0623100 clearly localize at the interface with the host cell during both the liver and blood stages of the infection and their expression and localization patterns so closely mimic those of IBIS1, we chose to further investigate the two tryptophan-rich domain-containing proteins. Because the current naming systems for genes containing this tryptophan-rich domain are inconsistent across species and none are ideal for indicating orthologs across species, we propose naming these two proteins based on their localization to intraerythrocytic *Plasmodium* induced structures (IPIS). We therefore refer to PBANKA_0524300 as IPIS2 and PBANKA_0623100 as IPIS3.

### Localization of putative *P. vivax* orthologs in *P. berghei*

Unlike IBIS1, which appears to be exclusive to the rodent *Plasmodium* species, tryptophan-rich domain-containing proteins are found across the *Plasmodium* genus (Figure S3). Syntenic orthologs of IPIS2 are conserved across the *Plasmodium* clade and are expressed in *P. vivax, P. knowlesi, P. malariae* and *P. cynomolgi*. Notably, no syntenic IPIS2 ortholog exists in the Laverania *Plasmodium* species *P. reichenowi* or *P. falciparum*. IPIS3 has syntenic orthologs only in therodent *Plasmodium* species; however, PVP01_0202200/*Pv*TRAg2 was previously annotated as an ortholog on PlasmoDB because of sequence similarity. In order to determine if the putative orthologs from *P. vivax* also targeted the membrane structures in the cytoplasm of *Plasmodium*-infected red blood cells, we generated *P. berghei* lines that expressed HA-tagged *Pv*TRAg8, the syntenic *P. vivax* ortholog of *IPIS2*, or HA-tagged *Pv*TRAg2, which shares similarity to IPIS3. These genes were integrated into the *P. berghei* genome such that they are expressed under the control of the *IPIS2* or *IPIS3* promoters, respectively. Both *Pv*TRAg8-HA and *Pv*TRAg2-HA are exported efficiently by *P. berghei* and localize to discrete structures in the cytoplasm of infected erythrocytes (Figure 3B).

### Absence of IPIS2 or IPIS3 confers a fitness disadvantage during *Plasmodium* growth in the blood

In order to determine if *IPIS2* or *IPIS3* contribute to successful growth of the parasite, each gene was deleted and replaced by selection markers via double homologous recombination (Figure S4). The successful gene deletion and selection of isogenic knockout lines indicated that neither IPIS2 nor IPIS3 are essential for blood-stage growth. Infection of *Anopheles stephensi* with the knockout lines and resulting development of salivary gland sporozoites revealed that these genes are also dispensable for *Plasmodium* development in mosquitoes (Figure S5A, B).

To assess the ability of the knockout lines to establish infection in mammals, C57BL/6 mice were infected with IPIS2- or IPIS3-deficient sporozoites. The wild-type and *ipis2*-*P. berghei* lines were detected in the blood between day three and four after infection with 1,000 sporozoites. In contrast, the *ipis3*-parasite line became visible in the bloodstream as late as five days after infection, and in two of the ten infected mice, IPIS3-deficient parasites were never detected in the blood (Figure 4A). This delay in patency prompted us to examine growth of the parasite lines in the livers of infected animals. Quantification of the parasite burden in the liver 42 hours after infection with sporozoites revealed no significant differences in growth of the *ipis2-* or *ipis3-* lines in the liver in comparison to wild-type *P. berghei* (Figure 4B). In parallel we assessed the liver-stage development of the knockout lines in cultured hepatoma cells *in vitro*. The size of the exo-erythrocytic forms in HepG2 cells were determined 48 hours after infection with sporozoites from each line, and no restriction of parasite growth was detected in the absence of IPIS2 or IPIS3 *in vitro* (Figure S5C), which is consistent with the *in vivo* data.

**Figure 4.**
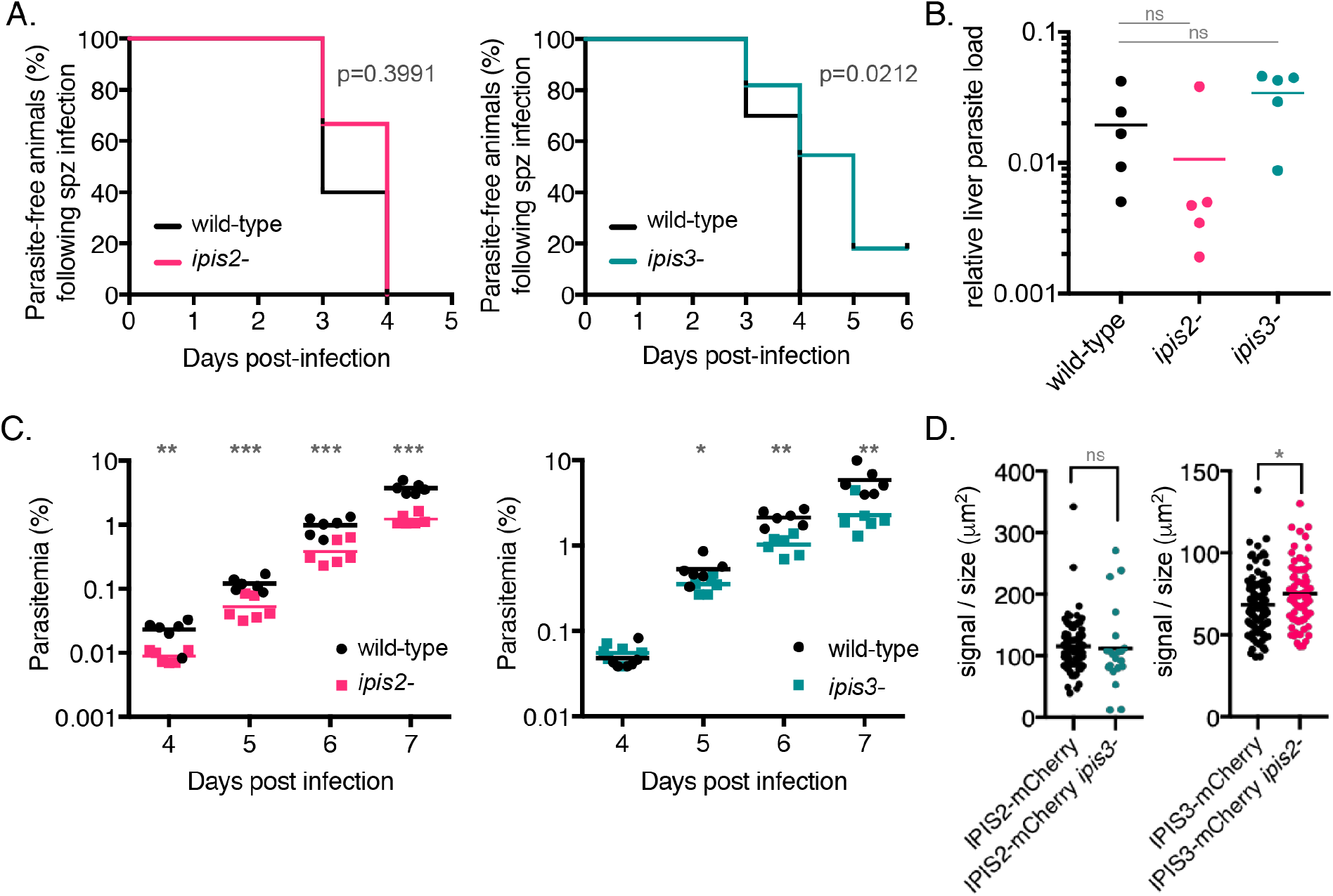
*ipis2-* and *ipis3-* contribute to blood-stage growth efficiency during *P. berghe*i infection. A) The appearance of *ipis3-* parasites in the blood following sporozoite injection is delayed in comparison to wild-type. C57BL/6 mice were infected with 1,000 sporozoites of either *ipis2-, ipis3-* or wild-type parasites, and blood-stage infections were monitored starting from day 3 post-infection with Giemsa-stained blood smears. Statistics: Log-rank (Mantel-cox) test; n=5 (*ipis2-*), n=10 (*ipis3-*). B) The growth of *ipis2-* and *ipis3-* parasites in the liver does not differ significantly from wild-type *P. berghei*. The relative parasite burden in the liver was measured by qPCR of RNA extracted from livers forty-two hours after intravenous injection of either *ipis2-, ipis3-*, or wild-type sporozoites. Levels of *Pb*18s rRNA were normalized to levels of mouse GAPDH RNA in each infected liver. Statistics: unpaired two-tailed t-test; n = 5; ns: not significant. C) The *ipis2-* and *ipis3-* lines do not grow as efficiently as wild-type in the blood of co-infected animals. Mice were infected with equal amounts of wild-type *P. berghei*-infected erythrocytes and *ipis2-*[mCherry; PyrS] or *ipis3-*[mCherry; PyrS] infected erythrocytes. Parasitemia was measured by flow cytometry between days 4-7 after infection. * p <0.05, ** p < 0.01, *** p < 0.005 determined by unpaired two-tailed t-test; n = 5. D) The absence of one IPIS protein does not dramatically influence the expression level of the other. Erythrocyte-associated mCherry signal was quantified in fixed unstained infected cells. The mCherry signal intensity was plotted as a ratio to parasite size to account for potential differences in expression in different stages. * p <0.05, unpaired two-tailed t-test.

Although the time to patency was statistically similar in mice infected with either wild-type or *ipis2*-sporozoites, the parasitemia of IPIS2-deficient parasites in the blood was on average lower than wild-type *P. berghei* following sporozoite infection (Figure S5D). The delay in patency observed upon infection with *ipis3-* sporozoites was accompanied by a corresponding decrease in parasitemia during the blood stages (Figure S5D). That *P. berghei* suffers growth consequences in the absence of IPIS2 or IPIS3 is underscored by the observation that mice infected with *ipis2*- or *ipis3*-sporozoites are less likely to succumb to infection within the first 12 days of infection (Figure S5E) in comparison to mice infected with wild-type *P. berghei*, which indicates reduced pathogenicity of the knockout lines.

Competitive growth analyses, in which mice are infected with both wild-type and knockout lines, have successfully identified growth phenotypes in the blood of infected mice [36, 37]. We selected for knockout parasites expressing single fluorescent markers as described [38], since the expression of distinct genetically-encoded fluorescent proteins in the wild-type and knockout lines allows their differentiation by flow cytometry. To assess growth of the knockout lines specifically in the blood stage of infection, mice were infected with equal numbers of red blood cells infected with wild-type and either *ipis2*- or *ipis3*-parasites. (Figure 4C). Although the same number of parasites were injected intravenously, fewer IPIS2-deficient parasites were detected in the blood of infected animals compared to wild-type parasites at each time point from four to seven days post-infection. While similar numbers of *ipis3*- and wild-type parasites were detected initially at day four, the wild-type line out-competed IPIS3-deficient parasites as the infection progressed. These results suggest that both knockout lines grow less efficiently in the blood than wild-type parasites.

The localization of IPIS2 and IPIS3 to the intraerythrocytic membrane structures and their requirement for efficient blood-stage growth prompted us to investigate whether the membrane structures are still formed in the cytoplasm of erythrocytes infected with the *ipis2*- and *ipis3*-knockout lines. In order to visualize these structures in the knockout lines, we tagged IPIS2 with the fluorescent protein mCherry in the IPIS3-deficient line, and likewise, tagged IPIS3 with mCherry in the line lacking IPIS2 (Figure S6). The IPIS proteins still appear in punctate structures distributed across the erythrocyte cytoplasm in the absence of either IPIS2 or IPIS3 (Figure S7A). Because IPIS2 and IPIS3 share expression and localization patterns in addition to the tryptophan-rich domain, we hypothesized that upregulation of one may compensate for the absence of the other. Because the fluorescent tags are on the natively expressed proteins, we were also able to use these lines to compare expression of IPIS2 and IPIS3 in the knockout lines based on the fluorescence intensity of mCherry. To account for the increase in expression of both IPIS2 and IPIS3 over the course of normal parasite development in red blood cells (Figure S7B), we normalized the fluorescence intensity of the IPIS2-mCherry or IPIS3-mCherry signal to parasite size in mixed blood stages. We detected no difference in the expression of IPIS2-mCherry or IPIS3-mCherry in the *ipis3-* or *ipis2*-lines, respectively, in comparison to the expression of the mCherry-tagged proteins in a wild-type genetic background (Figure 4D). These results indicate that IPIS2 and IPIS3 are not upregulated in the knockout lines.

### Schizonts are detected in the circulation in the absence of IPIS2 or IPIS3, but only IPIS2 is required for adherence of schizonts to CD36

Schizonts are only rarely detected in peripheral blood of *P. berghei*-infected mice due to their sequestration in lung and adipose tissue [39]. In animals infected with the IPIS2- or IPIS3-deficient lines, we detected schizonts in the peripheral blood during blood-stage growth more often than in animals infected with wild-type *P. berghei* lines (Figure S5F). Circulating schizonts can be recognized and cleared by the spleen, and infection of mice with *P. berghei* lines that fail to sequester out of circulation has been correlated with an increase in the weight of spleens of infected animals [17, 39-41]. In comparison to mice infected with wild-type *P. berghei*, the spleen-to-body weight ratio was on average 29% higher in mice infected with the ipis2-line and 19% higher in mice infected with the *ipis3*-line (Figure 5A). This data further indicated that sequestration may be compromised in red blood cells infected with the knockout parasites.

**Figure 5.**
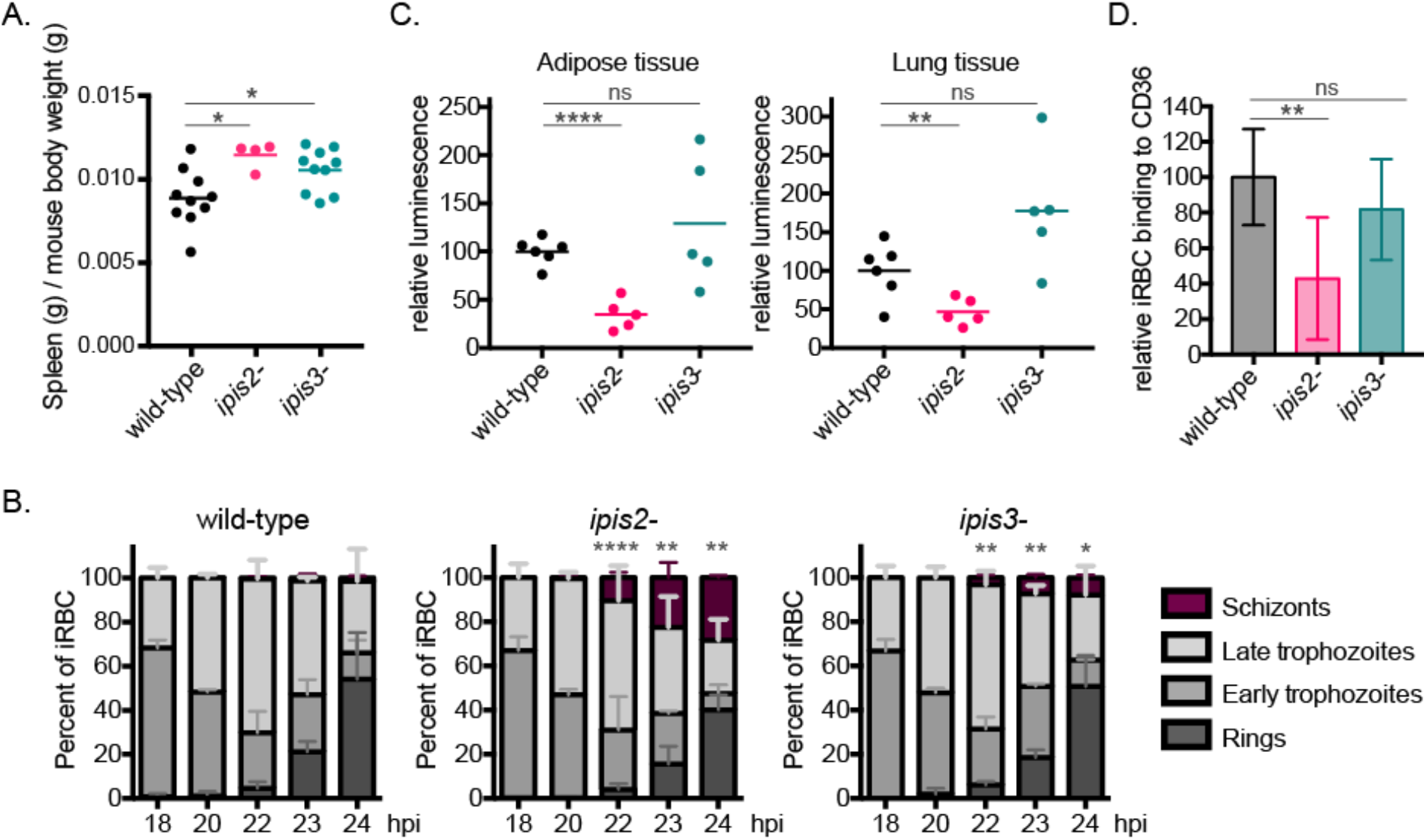
IPIS2 contributes to efficient sequestration of schizonts from the peripheral circulation. A) *ipis2-* or *ipis3-* infection increases spleen weight more than wild-type *P. berghei* infection. Mice were infected with sporozoites of either wild type (n = 10), *ipis2-*[GFP-Luc; PyrS] (n = 4), or *ipis3-*[GFP-Luc; PyrS] (n = 10) lines, and 8 days after infection the spleens were dissected and weighed. The spleen weights were plotted as ratios to mouse body weight.* p <0.05, unpaired two-tailed t-test. B) The absence of IPIS2 or IPIS3 resulted in increased schizonts in peripheral blood compared to wild-type *P. berghei* during synchronous infections. Mice were infected with *P. berghei* schizonts (wild-type, *ipis2-*[GFP-Luc; PyrS] or *ipis3-*[GFP-Luc; PyrS]) and the parasite stages detected in peripheral blood smears were quantified at the indicated time points following infection. n=5 mice in three experimental replicates. Statistical annotations (* p <0.05, ** p < 0.01, *** p < 0.005) refer to percent of schizonts in the transgenic line in comparison to the wild-type line at the indicated time points, unpaired two-tailed t-test. C) Tissue-associated parasite material was assessed by luminescence in homogenized tissues 23 hours after infection with *P. berghei* schizonts (wild-type *Pb* GFP-Luc, *ipis2-*[GFP-Luc; PyrS] or *ipis3-*[GFP-Luc; PyrS]). Values are normalized to organs from wild-type-infected mice.** p < 0.01, **** p < 0.0001, ns: not significant, unpaired two-tailed t-test. D) Red blood cells infected with IPIS2-deficient *P. berghei* do not bind efficiently to CD36-expressing CHO cells. Schizonts were incubated with CHO cells and the number of infected-red blood cells bound per cell quantified. The background binding to control CHO cells was subtracted, and the values were normalized to wild-type-infected red blood cell binding. The depicted data comprises three experimental replicates with three technical replicates for each parasite line and CHO cell line. ** p < 0.01, ns: not significant, unpaired two-tailed t-test.

In order to compare circulating schizonts directly and quantitatively across multiple lines, synchronized infections in mice were required. To this end, purified schizonts were injected intravenously into naïve animals, and the relative proportions of each life cycle stage present in blood smears taken between 18-24 hours after infection were quantified (Figure 5B). The injection of purified late-stage parasites resulted in a maximum of fifteen-fold more schizonts in the peripheral blood compared to wild type in the absence of IPIS2, and five-fold more schizonts in the absence of IPIS3. The parasitemia in animals 18-20 hours after injection of schizonts was similar across parasite lines (Figure S5G). The parasites used in this assay express GFP-Luciferase. Twenty-three hours after infection, the time when we saw the maximal number of schizonts, animals were sacrificed and the luciferase activity in the lung and adipose tissue was assessed. The luciferase activity in these tissues was lower in animals infected with the *ipis2*-line in comparison with wildtype-infected animals (Figure 5C). This result is consistent with the hypothesis that the wildtype schizonts sequester in these tissues more efficiently than the IPIS2-deficient line. In contrast, no significant difference in tissue-associated luciferase activity was seen with the *ipis3*-infected animals compared to wild-type parasites. Organs from these animals infected with IPIS3-deficient parasites had a wide range of luciferase activity, but the data suggest that there is no reduction in tissue-associated parasites caused by the absence of IPIS3.

*P. berghei* schizont sequestration in mouse tissues has shown to be mediated by binding to CD36 in the host endothelium [39]. To examine the ability of erythrocytes infected with *P. berghei* schizonts lacking IPIS2 or IPIS3 to bind CD36, blood infected with late blood stage parasites was incubated with CHO cells modified to express CD36 on the cell surface [42] (Figure 5D). Erythrocytes infected with wild-type *P. berghei* bound to the CD36-expressing cells on average five times better than to control CHO cells that do not express CD36. The adherence of erythrocytes containing IPIS2-deficient parasites to CD36 was reduced by about 60% compared to wild-type-infected cells. In contrast, there was no difference in binding of cells infected with the *ipis3*-line in comparison to wild-type *P. berghei*, indicating that erythrocytes infected with IPIS3-deficient parasites remain capable of adhering to CD36. Taken together, these data demonstrate that the absence of these exported IPIS proteins influences the blood infection stage. The absence of IPIS2 in particular influences the interaction between *P. berghei* schizonts and the host tissue endothelium, making IPIS2 required for efficient sequestration of schizonts from the blood circulation.

## Discussion

The formation of intracellular membrane structures containing parasite proteins is a conserved feature within the *Plasmodium* genus and occurs in all intracellular stages. These membranes demarcate regions where the parasite can interact with the environment of its host, and the functional activities occurring at these membranes will depend on their protein and lipid compositions.

The collection of proteins known to localize to the Maurer’s cleft-like structures in cells infected by *Plasmodium* species other than *P. falciparum* is limited, and this study has expanded this protein repertoire. By immunoprecipitating a *P. berghei* protein (IBIS1) present in host-localized membranes across intracellular stages, we identified additional proteins present in the IBIS in the cytoplasm of *P. berghei*-infected erythrocytes. Although neither IPIS2 nor IPIS3 contain a predicted PEXEL/HT (Plasmodium export element / host targeting) motif [43], they are exported from the parasite into the host erythrocyte where they localize to IBIS. We cannot determine from this data whether IPIS2 and IPIS3 interact with each other and/or IBIS1 directly or indirectly as part of one or more protein complexes. However, colocalization of the candidate proteins with IBIS1 suggests the IBIS1 pulldown successfully identified proteins present in the membrane structures formed in the host erythrocyte cytoplasm.

Previous knock-outs of IPIS2 showed no severe effect on blood stage growth [40], however large scale screening suggested there indeed was a slower growth phenotype associated with the knockout of IPIS2 and of IPIS3 [44]. We found the absence of IPIS2 influences the ability of *P. berghei* schizonts to sequester from the blood circulation. In support of previous findings [17, 40, 41, 45], efficient CD36-dependent cytoadherence supports robust *in vivo* growth, and we found that the failure of *ipis2*-schizonts to sequester from the circulation correlates with a fitness disadvantage during the blood infection stage.

The conclusion that IPIS2 is required for efficient sequestration of late-stage *P. berghei* from the circulation is supported by the detection of *ipis2*-schizonts in the peripheral blood and their relative absence from the tissues. At the end of one developmental cycle, wildtype schizonts are mostly absent from circulation and can instead be found in the lung and adipose tissue. Although *ipis3*-schizonts are found in the peripheral blood at this time, *ipis3*-parasite material can also be detected in the tissues of perfused animals. In contrast, the tissue-associated *ipis2*-parasite material is notably reduced, and this correlates with higher numbers of circulating *ipis2*-schizonts. Adherence assays using CD36-expressing cells demonstrated that erythrocytes infected specifically with IPIS2-deficient parasites fail to bind CD36 as efficiently as cells infected with wild-type *P. berghei*. The fitness cost of not sequestering is due to increased splenic clearance of infected cells [40], and we see a corresponding increase in spleen weight in animals infected with the *ipis2*-knockout line.

Deletion of the *IPIS3* gene also affected parasite fitness, but the phenotype appears to be distinct from that caused by the loss of *IPIS2*. The *ipis3*-line grows poorly in the blood in comparison to wild-type *P. berghei*; however, IPIS3-deficient schizonts are competent in CD36 binding and are tissue-associated in synchronized infections. The minor growth phenotype could therefore be due to a number of possible causes. Although ipis3-schizonts can still associate with tissues, a proportion of them can still be detected in the circulation. Synchronized growth suggested no major defect in the transition of *ipis3*-parasites from rings to schizonts, but protracted development in the late stages or defects in parasite exit from the red blood cell would increase the time *ipis3*-parasites spend as schizonts and perhaps consequently the likelihood that schizonts can be detected in circulation. These circulating schizonts may be sufficient to cause the increase in spleen weight that is observed in *ipis3*- -infected mice in comparison to wild-type infection. Alterations in parasite deformability or differences in how the *ipis3*-parasites trigger immune responses and inflammation may also be responsible for these observations.

The loss of IPIS3 also affects the transition between the liver and blood growth stages. Although *ipis3*-growth in the liver appeared to be robust when parasite burden was measured following sporozoite infection, we saw a delay in the time to patent blood-stage infection, and *ipis3*-parasites failed to establish blood-stage infection in a small proportion of infected mice. Comparisons to the *ipis2*-line indicate that the defect in *ipis3*-blood stage growth is not sufficient to cause this effect on patency. An intriguing possibility is that a defect in merozoite development or parasite egress in the liver and blood could explain the phenotypes of the *ipis3*-line, but these two phenotypes may also have alternative and independent causes.

Infected erythrocytes circulating freely in the blood represent only a proportion of the parasites in *Plasmodium*-infected vertebrates [46-50]. Although cytoadherence to the host vasculature and sequestration out of the blood circulation are most prominent in *P. falciparum* infections, these traits are not exclusive to this species. While some tissue-associated parasites are extravascular [50, 51], there is also evidence of intravascular sequestration of red blood cells infected with various *Plasmodium* species: Erythrocytes infected with *P. berghei* schizonts sequester in the tissue vasculature in a manner dependent on host CD36 [39]. Although late-stage *P. knowlesi* or *P. vivax* can be detected in the blood, erythrocytes infected with *P. knowlesi* can accumulate in the brain vasculature [52] and can bind to ICAM-1 or VCAM [53]. A proportion of late-stage *P. vivax* are sequestered out of circulation [46, 47, 50], and *P. vivax*-infected erythrocytes have been shown to bind to endothelial cells [47, 54-56]. *P. vivax* members of the PIR protein family appear to be involved in this binding [10, 54, 57]; however evidence that PIR proteins play a role in cytoadherence of other *Plasmodium* species where they are expressed is lacking [58, 59].

The molecules mediating the binding of *P. berghei*-infected erythrocytes to host CD36 are thus far unknown. Although the IPIS2 knockout line fails to bind to CD36 and sequester efficiently, the intracellular localization of IPIS2 precludes its direct involvement in cytoadherence. We therefore infer that there is an alteration in parasite-induced modification of the host cell surface in cells infected with the *ipis2*-line. The IBIS-localized proteins *Pb*SBP1 and *Pb*MAHRP1 have also been shown to be essential for efficient sequestration of *P. berghei* schizonts from the blood circulation [16]. Expression of IPIS3-mCherry in the *ipis2*-line confirmed that absence of IPIS2 causes no broad defect in parasite protein export or protein trafficking to IBIS, but failure of *Pb*MAHRP1 or *Pb*SBP1 to localize to IBIS in the *ipis2*-line would be expected to result in loss of schizont sequestration.

The requirement of *Pb*SBP1 and *Pb*MAHRP1 for *P. berghei* sequestration indicated that the IBIS formed in *P. berghei*-infected cells are functionally related to the Maurer’s clefts of *P. falciparum*-infected cells [17]. Although it is not yet known if the *P. knowlesi* ortholog of SBP1 (*Pk*SBP1) contributes to sequestration, *Pk*SBP1 has been found to localize to the Sinton and Mulligans Stippling – an encouraging suggestion for conserved protein roles in these membrane compartments [18]. While SBP1 and MAHRP1 were first described in *P. falciparum* [60, 61], IPIS2 and IPIS3 do not have definitive orthologs in *P. falciparum*. The gene encoding IPIS2 is, however, conserved and syntenic across the primate- and human-infecting *Plasmodium* clade, such as *P. vivax* and *P. knowlesi*. Our data consequently support the idea that membrane structures formed in the infected erythrocyte cytoplasm play conserved roles in the host cell remodeling that facilitates the interactions between the infected cell and the host tissues.

Both IPIS2 and IPIS3 possess a tryptophan-threonine-rich domain (PFAM: PF12319) that is found in multiple *Plasmodium* proteins. Proteins with this domain are sometimes referred to as TRAg, PyAg or Pv-fam-a. Importantly however, in *P. berghei* Fam-A refers to an unrelated family of proteins, which contain a predicted START domain. The tryptophan-threonine-rich domain is characterized by positionally conserved tryptophan residues, and proteins containing this domain can be immunogenic and are implicated in a range of processes including adherence to and invasion of erythrocytes [62-65]. Our study shows for the first time that a protein with this tryptophan-rich domain is involved in remodeling of the host erythrocyte and supports the idea that this domain does not necessarily define the function of these proteins. The family of proteins containing this domain is expanded in the *Plasmodium* clade of *Plasmodium* species, which contains all human-infecting species apart from *P. falciparum*. For example, *P. vivax* encodes forty Trp-Thr-rich domain-containing proteins and *P. knowlesi* has twenty-nine. In contrast, *P. falciparum* encodes only three proteins with this domain. Various expression and localization patterns have been described for *Plasmodium* proteins possessing this domain [64, 66]. A *P*.*falciparum* member of this family, TryThrA, was recently shown to immunoprecipitate with *Pf*SBP1 and localize to Maurer’s clefts [67]. However, genetic targeting of TryThrA did not influence the ability of *P. falciparum*-infected erythrocytes to cytoadhere to CD36. In this study, we demonstrate that putative *P. vivax* orthologs of IPIS2 and IPIS3 appear to engage the protein export machinery when expressed in *P. berghei* and, similar to *Pb*IPIS2 and *Pb*IPIS3, localize to punctate structures in the cytoplasm of *P. berghei*-infected erythrocytes. When the *P. berghei* protein IBIS1 was expressed in *P. falciparum*, it was found to localize to the Maurer’s clefts [68], which further supports the cross-species export recognition we also demonstrate here with the expression of *P. vivax* proteins in *P. berghei*. While the results of trans-species expression should be interpreted with caution, these data may suggest that these orthologs may have a similar target in the cytoplasm of *P. vivax*-infected red blood cells.

In addition to colocalizing with IBIS1 in infected erythrocytes, IPIS2 and IPIS3 are also expressed in the liver stage, and like IBIS1, they surround the liver-stage parasites. Such dual expression and differing localizations in these two stages have also been demonstrated for multiple other exported *Plasmodium* proteins [58, 69]. *P. berghei* members of the PIR, Fam-A and Fam-B protein families have been shown to be expressed in both stages, exported into the host erythrocyte and retained in the liver-stage parasitophorous vacuole [58]. It should be noted that in these examples, proteins were tagged with the fluorescent protein mCherry, as was done here with IPIS2 and IPIS3. While fluorescent protein tags do not hinder protein export across the parasitophorous vacuole in blood stages, we cannot exclude the possibility that this tag may interfere with protein translocation in the liver stage of infection. Although the localization differs in the two infection stages, IPIS2 and IPIS3 are at the boundary with the host cell in both, which is noteworthy considering the physiological differences between erythrocytes and hepatocytes.

In parallel to our IBIS1-mCherry pulldown, we immunoprecipitated mCherry-tagged EXP2 from *P. berghei*-infected erythrocytes. We not only pulled out previously characterized components of the PTEX apparatus, which validated our experimental approach and analysis, but we also identified and highlighted some additional candidate proteins that have been previously immunoprecipitated with HA-tagged PTEX components from *P. falciparum*-infected cells [33, 34]. These proteins, marked in light blue in Figure 1, were pulled down previously with either PTEX150 or PTEX88 [33, 34] and have not yet been investigated, but their association with different PTEX components in multiple *Plasmodium* species makes them strong candidates for having functional association with the PTEX complex in infected cells. Nevertheless, these and the other novel proteins we found associating specifically with EXP2 need further characterization before any role in protein export can be established. Likewise, our validation of IPIS2 and IPIS3 as resident proteins of the IBIS in infected erythrocytes suggests that the additional proteins we detected specifically in the IBIS1-mCherry immunoprecipitates are potential protein components of the cytoplasmic membrane structures of *Plasmodium*-infected erythrocytes.

Multiple *Plasmodium* species have cytoadhesive potential, and parasite sequestration has broad consequences for disease progression and parasite persistence [70-73]. Studying both the conserved and the species-specific features that mediate host cell remodeling can improve our understanding of how *Plasmodium* species promote favorable interactions within their hosts’ tissues. Our studies show IPIS2 and IPIS3 localize to erythrocyte-localized membrane structures in *P. berghei*-infected cells and suggest that orthologs of these proteins may localize to such membranes across several of the *Plasmodium* species in which protein export and host remodeling are less well characterized. Our results support the idea that the membrane structures in the cytoplasm of *Plasmodium*-infected cells have conserved functions across species. While variable in morphology, they are important for the remodeling that allow *Plasmodium*-infected cells to navigate the tissues of their host.

## Materials and Methods

### Ethics statement

Animal research was performed in strict adherence to the “Tierschutzgesetz in der Fassung vom 22. Juli 2009”, which implements the Directive 2010/63/EU of the European Parliament and Council “On the protection of animals used for scientific purposes.” The protocol for usage of animals obtained approval from the Berlin ethics committee (Landesamt für Gesundheit und Soziales Berlin) and operated under the permit numbers G0469/09 and G0294/15.

### Experimental animals, cells and parasite lines

Female NMRI and C57BL/6 mice were either purchased from Charles River Laboratories (Sulzfeld, Germany) or bred in-house. NMRI mice were used for blood-stage growth assays. Sporozoite inoculations, spleen weight measurements, and sequestration experiments were performed using C57BL/6 mice. The *P. berghei* reference lines used were *P. berghei* ANKA, *P. berghei* GFP-Luc (RMgm-29) [74], and *Plasmodium berghei* Bergreen [75]. The HepG2 and Huh7 hepatoma cell lines were cultured in DMEM supplemented with fetal calf serum (FCS). Chinese hamster ovary (CHO) cells expressing GFP or CD36-GFP were a gift from the laboratory of Iris Bruchhaus (BNITM, Hamburg) [42] and were maintained in Ham’s F12 supplemented with 10% FCS and 1 mg/ml G418. *Anopheles stephensi* mosquitoes were raised under a 14 h light/10 h dark cycle at 28°C and 80% humidity and were fed daily on 10% sucrose.

### Construction of plasmids

For the construction of the plasmids used for endogenous C-terminal mCherry tagging, regions containing the *IPIS2, IPIS3* or *PBANKA_1400700* genes were amplified from *P. berghei* genomic DNA using the respective primers (Table S2) and integrated into the SacII and XbaI sites of the B3D+mCherry plasmid [76]. The sequences of *Pv*TRAg8 and *Pv*TRAg2 were codon optimized (sequences provided in Figure S8), synthesized (GenScript) and cloned into the NotI and SpeI sites of b3D.DT^H.^D (provided by Dr. Andrew Waters, Glasgow University). The 5’ and 3’ untranslated regions of the *IPIS2* and *IPIS3* genes were cloned into the GOMO-GFP luciferase vector using the respective primers (Table S2) containing SacII and NotI sites and the XhoI and KpnI sites, respectively, in order delete *IPIS2* and *IPIS3* via double homologous recombination.

### Generation of transgenic *P. berghei* parasites

Transfection *P. berghei* schizonts and selection of transgenic parasites was carried out as previously described [74]. Briefly, the plasmid constructs used for genetic alteration were either linearized, or in the case of the knockouts, the vectors were digested with restriction enzymes whose cut sites flanked the disruption element. Schizonts were isolated from overnight cultures, and transfection of the schizonts was done using the Nucleofactor T-cell transfection kit in combination with an Amaxa gene pulser using the program U-33. Transfected parasites were injected into recipient mice, and after the second day, transgenic parasites were selected for using 70ng/μl, pH 3.5-5.5 pyrimethamine drinking water.

Clonal lines of *ipis2*-[GFP-Luc;mCherry], *ipis3*-[GFP-Luc;mCherry], and *ipis2*-*ipis3-* [GFP-Luc;mCherry] were generated by FACS sorting mCherry+GFP+ parasites and injecting them into new recipient mice. PCR was used to confirm genotypes (Table S2).

To remove the DHFR-resistance from the knockout lines and concurrently select for parasites encoding single fluorescent reporters [38], the knockout lines were put under selective pressure for five days with 1.5mg/ml 5-fluorocytosine (5-FC) to the mouse drinking water, which was administered fresh daily and protected from light. Isogenic clonal lines were generated by FACS sorting parasites that expressed only mCherry or GFP-luciferase. This process yielded the lines *ipis2*-[GFP-Luc; PyrS] and *ipis3*-[GFP-Luc; PyrS]. Genotypes were confirmed with PCR (Table S2). The lines *ipis2*-IPIS3-mCherry[GFP-Luc; PyrS] and *ipis3*-IPIS2-mCherry[GFP-Luc] were generated using *ipis2*-[GFP-Luc; PyrS] and *ipis3*-[GFP-Luc; PyrS], respectively.

### Cross fertilization of transgenic *P. berghei* lines

To generate parasites expressing both IBIS1-GFP and the mCherry-tagged candidate proteins, the lines were genetically crossed. Mice were co-infected with equal numbers of the mCherry-expressing *P. berghei* lines and the IBIS1-GFP *P. berghei* line and fed to *Anopheles stephensi* mosquitoes. Sporozoites comprising a mixed population including double-fluorescent parasites were isolated from the salivary glands of mosquitoes infected with cross-fertilized lines on day 21 after transmission to the mosquitos.

### Overnight *ex vivo* culture and purification of *P. berghei* schizonts

*P. berghei* schizonts were cultured *ex vivo* as described [74] in DMEM culture medium supplemented with FCS and heparin for up to 22 hours at 37°C on a shaking platform rotating at 15 rpm with a gas composition of 5% CO_2,_ 10% O_2_, and 85% N_2_. Schizonts were isolated from the culture through a one-step Nycodenz gradient separation using 55% Nycodenz in PBS.

### Immunoprecipitation of IBIS1-mCherry

Parasites were cultured as described above for 16-20 hours and late-stage parasites were purified with a 60% Nycodenz gradient. Cells were lysed on ice for thirty minutes with periodic agitation in a buffer containing 10 mM Tris/Cl pH 7.5, 150 mM NaCl, 0.5 mM EDTA and 0.5 % NP40 supplemented with 1mM PMSF and a protease inhibitor cocktail (Sigma-Aldrich). Immunoprecipitations from cell lysates were performed with the RFP-Trap-A kit (Cromotek) according to manufacturer’s instructions using equal numbers of parasites per line, which varied from 7×10^8^ to 7×10^9^ across experimental replicates. The *P. berghei* lines used included a line expressing IBIS1-mCherry [15], a line expressing mCherry from the HSP70 promoter [28], a line in which mCherry is fused to the predicted PEXEL motif of the circumsporozoite protein and expressed from the IBIS1 promoter [29], and a line expressing EXP2-mCherry [30].

Proteins eluted from the IP beads were digested using an automated sample-preparation workflow (Axel-Semrau Proteome Digest-O-r [77]. Briefly, the samples were reduced by 1 mM tris(2-carboxyethyl)phosphine (TCEP, Merck) and free sulfhydryl groups carbamidomethylated using 5.5 mM chloroacetamide (Sigma-Aldrich). Proteins were digested with 0.5 µg sequencing grade endopeptidase LysC (Wako) overnight at room temperature. The reaction was terminated by adding trifluoroacetic acid (TFA, Merck) to a final concentration of 1% resulting in a final pH of 2. The peptides were purified using C18 stage-tips (Empore SPE disks, 3M) [78] and measured on a Q-Exactive plus mass spectrometer (Thermo-Fisher, Germany) coupled to a nano-LC system (easy-nLC, Thermo-Fisher, Germany). An amount of 2 μg of the peptide sample were injected and separated using a 3h gradient (4 to 76 % acetonitrile and 0.1 % formic acid in water) at a flow rate of 0.25 μl/min on an in-house prepared nano-LC column (0.075 mm x 250 mm, 3 µm Reprosil C18, Dr. Maisch GmbH). The separated peptides were ionized on a proxeon ion source and directly sprayed into the mass spectrometer (Q-Exactive Plus, Thermo). The MS1 acquisition was performed at a resolution of 70,000 in the scan range from 300 to 1700 m/z. The top 10 intense masses were selected for MS2 analysis. MS2 scans were carried out at a resolution of 15,500 with the isolation window of 2.0 m/z. Dynamic exclusion was set to 30 s and the normalized collision energy to 26 eV. For the automatic interpretation of the recorded spectral data, the MaxQuant software package version 1.6.0.16 was used [79]. Carbamidomethylation was set as a fixed modification while oxidized methionine and acetylated lysine were set as variable modifications. An FDR of 1 % was applied to peptide and protein level, and an Andromeda-based search was performed using a p. berghei Uniprot database.

The mean LFQ intensities across all three replicates were used for analyses. Enrichment in the experimental samples was determined by calculating the ratios of the intensities of all protein groups detected in the experimental samples relative to the control samples, and the binary logarithm of these ratios was formed. The standard normal distribution was calculated using the means and standard deviation for the obtained log2 ratios (Table S1).

### Blood stage immunofluorescence assays

For infected erythrocytes, 1µl of blood from an infected mouse was washed, diluted in PBS, and added to concanavalin A-coated coverslips (0.3mg/ml concanavalin A). The sample was then fixed with 4% PFA + 0.0075% glutaraldehyde in PBS for 15 minutes at room temperature.

Fixed cells were permeabilized with 0.1% triton X-100 in 3% BSA/ PBS and then blocked with a 3% BSA/ PBS for 1 hour. The samples were subsequently probed with antibodies against RFP (5F8; Chromotek; 1:500) and GFP (ab13970; Abcam; 1:1000).

### Calculation of mCherry signal in infected blood

Infected erythrocytes were fixed and mounted in DAPI-containing Fluoromount-G (SouthernBiotech). Fluorescence microscopy was done using an Axio Observer Z2 epifluorescence microscope (Zeiss) and 63x/1.4 objective. Images were taken with AxioCam MR3 camera at identical exposures to quantify the mCherry signal in the presence and absence of *IPIS2* and *IPIS3*. Images were imported into FIJI for quantification of both the mCherry signal in the red channel and parasite size in the DIC channel [80].

The outline of the red blood cell visible in the DIC channel was used to define the area of mCherry calculation within the red channel. To account for the asynchronous infections, parasite size was used as a proxy for parasite development, and the mCherry signal intensity calculated was expressed as a ratio to parasite size, which was also determined in the DIC channel.

### Infections with *P. berghei* sporozoites

Sporozoites were extracted from salivary glands of infected mosquitoes, and 1,000 sporozoites in RPMI were injected into the tail vein of naïve mice for assessing growth, and 10,000 sporozoites were injected for qPCR to determine the parasite burden in the infected liver. Thin Giemsa-stained blood smears were made every day starting from day 3 after injection in order to determine parasitemia by light microscopy. To assess the weight of spleens in infected animals, eight days after the injection of 1,000 sporozoites, the infected mice were weighed and sacrificed. The spleens were then dissected and weighed. To account for differences in body weight, a ratio of spleen weight to body size was calculated.

For *in vitro* infections, HepG2 cells were seeded and infected with isolated sporozoites. After addition of sporozoites, cells were centrifuged at 1800 x *g* for 5 minutes. The cells were fixed with 4% PFA for 15 minutes. The fixed cells were permeabilized with methanol and then treated with a block + quenching solution of 3% BSA + 4% FCS serum in ammonium chloride in PBS for 1 hour at RT.

Antibodies against RFP (ab62341; Abcam), GFP (ab13970; Abcam) and *Pb*HSP70 [81] were used for immunofluorescence analyses. All prepared samples were visualized with fluorescence microscopy. Exoerythrocytic form size of the developing parasites was calculated using FIJI imaging software [80].

### qPCR determination of parasite burden in infected mouse livers

The parasite burden in infected mouse livers was determined with qPCR quantification of parasite 18s rRNA. Mice were infected i.v. with 10,000 sporozoites. 42 hours post infection, the mice were sacrificed, and the livers harvested. They were washed briefly in PBS before homogenization in TRIzol. Liver homogenates were diluted 1:1 in TRIzol and stored at -80°C. mRNA was extracted using the Direct-zol RNA MiniPrep Plus kit according to the manufacturer’s protocol. The extracted mRNA was diluted to 2ng/µl. cDNA was generated using RETROscript (Invitrogen) according to the manufacturer’s protocol. Relative parasite burden was determined via quantitative PCR by comparing the mean Ct value of the *P. berghei* 18s ribosomal subunit (Gene ID: 160641) to the mean Ct value of the *Mus musculus* gapdh (Gene ID: 281199965) in the generated cDNA.

### Flow cytometry analysis of *P. berghei* blood stage infections

Flow cytometric detection of blood stage parasites was performed as described [36]. Blood stage co-infections were initiated with parasites transferred from donor mice in which the parasitemia was below 1% and quantified by flow cytometry on a BD FACS Canto II. The gating strategy was as described previously [36]. The blood from the donor mice was serially diluted such that 500 parasites of both wild type and either *ipis2-*[mCherry; PyrS] or *ipis3-*[mCherry; PyrS] in 100ul of RPMI were injected per animal into naïve recipient mice. On days 4-7 following injection, parasitemia of both lines was measured by flow cytometry from a drop of tail blood diluted in Alsever’s solution. Blood cells infected with parasites of both lines (the mCherry+GFP+ population) were excluded from the parasitemia calculation.

### Synchronous blood-stage infections

In order to assess schizont sequestration in a synchronous infection, schizonts were isolated from an overnight *ex vivo* culture as described above. Each mouse was injected with approximately 5×10^7^ infected red blood cells isolated from the overnight cultures. Giemsa-stained thin blood smears using tail blood were made at 2 hours post infection to ensure successful invasion of merozoites. After this, giemsa smears were made at the indicated time points after infection. The stages were identified by light microscopy as rings, early trophozoites, late trophozoites, and schizonts. Early trophozoites were classified when trophozoites were 1/3 the size of the red blood cell or smaller, and late trophozoites were classified when parasites were larger than 1/3 of the infected cell. 300 total parasites were counted per time point per mouse, and the population percentages were calculated as a portion of the total infected erythrocytes. For quantification of tissue-associated luciferase signal, mice were sacrificed and perfused 23 hours after injection of schizonts. The lung and adipose tissue were weighed and homogenized through a 70 µm cell strainer in RPMI medium and spun down at 1500 rpm for 3 minutes. The pellet was resuspended in 1 ml RPMI. The cell suspension was mixed 1:1 with ONE-GloTM Luciferse Assay Buffer (Promega), and the luminescence was measured with the Synergy HTX Multi-Mode Reader. To account for differences in organ weight, the ratio of tissue mass to luminescence was calculated. To include data from three experimental replicates, values were normalized to those obtained for the organs of the wild-type-infected mice.

### CD36 adherence assay

Experiments assessing the adherence of infected red blood cells to CHO cells expressing either GFP or CD36 were performed as previously described [42, 82]. 1×10^5^ CHO cells were seeded on coverslips in 24 well dishes. To obtain late-stage *P. berghei*, infected blood was incubated in ex vivo cultures as described above. At 16-18 hours blood was resuspended in binding media (RPMI, 2% glucose) and added to the CHO cells at 1.5×10^6^ infected cells per well. Cells were incubated at 37°C for 1 hour with periodic shaking, washed in binding media, fixed in 1% glutaraldehyde and stained with a 1:10 Giemsa-Weiser buffer solution. The number of infected red blood cells bound per CHO cell was assessed by microscopy. At least 250 CHO cells per coverslip were quantified. For each parasite line, the binding to the control GFP-expressing CHO cell line was assessed for reference. The experiment was performed three times with triplicates of all samples. To include data from all experiments, the binding was normalized to the binding of the wild-type *P. berghei*-infected cells

## Acknowledgements

We would like to thank Kai Matuschewski, Alex Maier and Andreas Herrmann for fruitful discussions, support and helpful feedback. We thank Nahla Galal Metwally and Iris Bruchhaus for sharing reagents and technical expertise. We furthermore thank Manuel Rauch, Carolin Rauch, Berit Söhl-Kielczynski and Werner Stenzel for technical assistance and Mariana de Niz for experimental advice. The authors acknowledge the instruments and expertise of Microscopy Australia at the Centre for Advanced Microscopy, Australian National University, a facility that is funded by the University and the Federal Government through NCRIS. This work was supported by the German Research Foundation (DFG) and the International research training group IRTG2290.

